# Natural Product Screen Identifies Asiatic Acid Targeting Mutant p53

**DOI:** 10.1101/2023.07.23.550189

**Authors:** Geeta Swargiary, Qazi S Jamal, Shreesh Ojha, Niraj Kumar Jha, Keshav K Singh, Shalini Mani

## Abstract

Mutations in p53 are common in different cancer types and are reported to protect cancer via different mechanisms. R175H, R248Q, and R273H are the hotspot mutations of p53 and are suggested to increase aerobic glycolysis in cancer cells. Cancer cells rely mostly upon aerobic glycolysis, so it will be interesting to target these three p53 mutants for designing alternative cancer therapy. The class of compounds studied for their potential to target energy metabolism of cancer cells is called mitocans. The current study is an approach to explore if these selected 3 mutants of p53 may act as suitable target(s) for natural mitocans. Hereby, we selected 60 phytocompounds altogether from *Andrographis paniculata* and *Centella asiatica* and docked against all three p53 mutants, using Autodock vina. 11 compounds were sorted based on their binding energies and drug-like properties, and toxicity levels prediction showed asiatic acid to be the most significant. As asiatic acid was observed to significantly bind with R248Q only, R248Q-Asiatic acid was identified for molecular dynamics simulation using GROMACS which showed significant interactions. In conclusion, mutant R248Q was observed to be the best target for asiatic acid, though additional *in-vitro* experiments are important to validate the findings of this study.

## 1. INTRODUCTION

*TP53* is a tumor-suppressor gene and encodes for a protein, present in the nucleus of cells. p53 is a transcription factor and exhibits a significant function in the regulation of cell division as well as cell death. In normal circumstances, p53 expression is very low, which is primarily due to its proteasomal degradation facilitated by E3 ubiquitin ligase MDM2 (Honda et al., 1997). In case of DNA damage, it undergoes post-translational modifications and is made to gather in the cell nucleus. These chemical modifications trigger the separation of MDM2 from p53 (Vousden et al., 2002; Lacroix et al., 2006; Prives et al., 1999). Functionally dynamic p53 transactivates a suitable number of its target genes, resulting in triggering cell cycle arrest followed by repairing of DNA and/or apoptotic cell death (Lacroix et al., 2006; El-Deiry, 2003). However, in case of severe DNA impairment, p53 employs its pro-apoptotic effects and thus abolishes the damaged cells. Hence, p53 has the capacity to sustain genomic integrity.

Being one of the most important transcription factors, p53 regulates the expression of different genes, important for metabolic pathways. For instance, p53 upsurges the level of cytochrome c oxidase 2 (SCO2), p53 modulator of apoptosis (PUMA), TP53-induced glycolysis and apoptosis regulator (TIGAR), glucose transporter 1 and 4 (GLUT1, GLUT4) (Puzio-Kuter, 2011) and glutaminase 2 (GLS2) but declines the level of phosphoglycerate mutase (PGM) in damaged cells. Interestingly, there is mounting evidence indicating that p53 is a vital performer in controlling cellular energy metabolism too (Kruiswijk et al., 2015; Berkers et al., 2013; Matoba et al., 2006). During oxidative stress, p53 activates the transcription of the *TIGAR*, reducing the amount of intracellular fructose-2,6-bisphosphate and then subsequently lowering the glycolysis rate as well as intracellular reactive oxygen species (ROS). Reduced glycolysis makes it possible for entering to pentose phosphate pathway (PPP), hence lowering the amount of cell death. In contrast to TIGAR, the level of PGM is decreased due to p53 activation, eventually reducing the rate of glycolysis.

*TP53* is the gene commonly found to carry mutations in cancer patients, and the mutated p53 is often addressed as the guardian of cancer cells (Mantovani et al., 2010). Human malignancies frequently harbor missense mutations in *TP53*, causing the loss of their tumor-suppressive properties. But the mutants apply trans-dominant suppression above their wild-type counterparts. The mutant p53, therefore, defends the cancer cells through this action. The mutant p53 not only loses its tumor suppressor role but acquires a new tumour-promoting role also called the gain of function (GOF) which drives metabolic rewiring in cancer cells, one of the crucial contributors to cancer progression (Liu et al., 2019; Hashimoto et al., 2019). According to the National Cancer Institute, GDC Data Portal, a total number of 1308 mutations of p53 are responsible for 4,703 cancer cases. Some of the most frequent mutations of p53 are R175H, R248Q, R273C, R273H, R248W, etc.

The potential of mutant p53 to maintain tumor cell survival was found to depend critically on its cytoplasmic localization and certain p53 mutants have been found to reside in mitochondria too (Mantovani et al., 2017). For example, the R175H and R273H mutations in *TP53* induce the mitochondrial citrate transporter protein (CTP), an integral membrane protein found in the mitochondrial inner membrane (MIM) of the, and help in the bidirectional shuttling of citrate between the mitochondria and cytosol. The shuttling further stimulates respiration and helps in maintaining the MIM integrity, thus not allowing the discharge of apoptotic molecules, like cytochrome c, and more ATP generation, promoting tumorigenesis (Blandino et al., 2020).

Amongst all the reported mutations of p53, R175H, R248Q, and R273H, are observed to be the most important mutations, responsible for influencing the pathways involved in energy metabolism by driving the Warburg’s effect (Zhang et al., 2013). These 3 mutations are noted to cause GLUT1 translocation to the cellular membrane by boosting the level of RhoA and ROCK, two small GTPases. RhoA and ROCK promote the Warburg effect and cancer cells favour aerobic glycolysis over oxidative phosphorylation (OXPHOS) as a means of energy production (Chiang et al., 2021). Furthermore, research revealed that in head and neck cancer, the mutants R282W, G245C, R175H, and P151S relocate to cytoplasm and limit the phosphorylation as well as the commencement of AMPK, a sensor of energy and have the ability to suppress aerobic glycolysis. As a result, mutant p53s suppress AMPK phosphorylation to cause the Warburg effect, which promotes tumor growth (Zhou et al., 2014). Thus, the association of mutant p53 (R175H, R248Q, and R273H) and energy metabolism pathways indicates the significance of targeting these three important p53 mutations for designing alternative cancer therapy.

As per the literature, a large number of natural/synthetic compounds (named Mitocans) are being studied for their ability to affect the bioenergetic of cancer cells and thus aiming to target their mitochondria (directly/indirectly). Based on different targets (such as enzymes of glycolysis, mitochondria DNA, respiratory chain enzymes, mitochondrial membrane proteins etc.), these mitocans are divided into different classes (Mani et al., 2020). Recently, we also explored the mitocan potential of the phytocompounds of *Centella Asiatica* (CA) and *Andrographis paniculata* (AP) against hexokinase 2 (commonly studied target for class I mitocans) and as a result, andrographolide, asiatic acid, and bayogenin were predicted to be the potential mitocans against HK2 (Swargiary and Mani, 2021). Following this study, we are further interested in investigating the mitocan ability of the same set of compounds against mutant p53 (R175H, R248Q, and R273H), as these 3 mutants have never been explored as possible targets for mitocans. Studying the mutant p53 as the target of mitocan(s) may help in designing the alternative/combinatorial therapy for managing the patients, carrying such mutations.

## 2. MATERIALS AND METHODS

### 2.1. Preparation of the Ligands and the Receptors

The 3D structures of all the 60 natural compounds of CA & AP from the Indian Medicinal Plants, Phytochemistry And Therapeutics (IMPPAT) database were retrieved. Similarly, the 3D structures for the selected mutant p53 proteins (R175H, R248Q, and R273H) are not available in the RCSB protein databank (https://www.rcsb.org). Thus, their structures were generated by the method of comparative modeling using the Modeller software. Initially, the wild-type p53 (P53WT) sequence was retrieved from the UniProt database (Entry ID: P04637, Entry Name: P53_HUMAN), and the mutations R175H, R248Q, and R273H were individually incorporated into their respective positions. Since only a single mutation was incorporated into the entire sequence, hence, only the P53WT structure with PDB: 2AHI, 1.85 Å resolutions was selected as the template. The generated models of P53WT and the three mutant p53s were validated by different applications of SAVES that include the ERRAT, VERIFY3D, and PROCHECK. Additionally, the generated model structures were superimposed against the template to understand the extent of deviation between the model and the template. The superimposition was performed by using the tool PyMOL.

### 2.2. Binding Pocket Identification and Molecular Docking

The modeled 3D structures were used for further molecular docking. Firstly, the binding pocket was identified from the recent report that suggested a novel drugable pocket in the DNA-binding region of the mutant p53. This pocket is defined by the following amino acid residues: Arg174, Glu192, Asp207, Asp208, Arg209, Asn210, Thr211, Phe212, Arg213, and His214 (Pradhan et al., 2019). By using the Autodock Tools, a grid box dimension of 20 Å × 20 Å × 20 Å was centered on the identified pocket residues to confine the search space of each docked natural compound. Molecular docking of the chosen ligands and the receptors was performed on Autodock Vina and their binding energies along with the ligand confirmations were noted. The natural compounds were further eliminated on the basis of their binding energies. The compounds with the minimum binding energies were chosen and the conformation of each ligand for the receptors after the docking was assessed on PyMOL and LigPlot+ to visualize the hydrogen as well as the hydrophobic bonds between the selected ligands and the receptors.

### 2.3. Druglikeness and Toxicity Predictions

Drug-like characteristics of the chosen 11 phytocompounds from the molecular docking analysis were predicted by using the online tool Molinspiration. The molecular properties comprising the molecular weight (MW), volume (vol), topological polar surface area (TPSA), miLogP, number of rotatable bonds (nrotb), atoms (natoms), hydrogen acceptors, and donors were provided by the Molinspiration. Further, the phytocompounds were eliminated based on the criteria adapted from Swargiary & Mani, 2021 which used the collective criteria of Veber’s rule, Ghosh’s rule, Pfizer’s rule, and Lipinski’s Rule of five (Swargiary & Mani, 2021). Moreover, the bioactivity score of the remaining phytocompounds after the elimination was also predicted by Molinspiration. The bioactivity score is predicted based on the very important drug targets containing the GPCR ligands, kinase inhibitors, ion channel modulators, nuclear receptors, protease inhibitors, and enzyme inhibitors.

ProTox-II web server, which is available for free, was used to calculate acute toxicity, organ toxicity (hepatotoxicity), and toxicity endpoints such as carcinogenicity, immunotoxicity, mutagenicity, and cytotoxicity. To calculate the toxicity of a small molecule, the ProTox-II online server accepts data in the form of a 2D chemical structure or SMILES notations and presents an overall toxicity class and radar chart for the given small molecule (Banerjee et al., 2018).

### 2.4. Molecular Dynamics Simulations (MDS) of the Docked Complex

After the collective analysis of molecular docking, drug-likeness predictions, and bioactivity scores the selected docked complex was subjected to MDS to understand the molecular interactions of the ligand and the receptor. Thus, the MDS environment was commanded to perform 20 ns simulations using GROningen MAchine for Chemical Simulations (GROMACS) tool 2018 version (Van Der Spoel et al., 2005). Simulations of P53WT and R248Q in water were achieved by GROMACS standard protocol. Initially, the topology file for the receptor was generated by pdb2gmx module of the GROMACS and then the CHARMM27 all-atom force field was selected. Any protein atom was solvated by simple point charge (spc) water molecules and was at least 1 nm distant from the box’s wall under periodic boundary conditions. Next, the ligand (AA) topology file was produced by the SWISSPARAM server (Zoete et al., 2011). To meet the need for electro-neutrality, NaCl counter ions were introduced, and energy minimized by the steepest descent approach. To maintain the system in a stable environment (300 K, 1 bar), Parinello-Rahman pressure coupling and Berendsen temperature coupling were utilized. The coupling constants for temperature and pressure were adjusted to 0.1 and 2.0 ps, respectively. The Coulomb cut-off (r coulomb) and neighbor list (rlist) were fixed at 0.9 nm, and the short-range Van der Waals interaction cut-off distance (rvdw) was set at 1.4 nm using the partial mesh Ewald (PME) technique. The LINCS approach (Duan et al., 2014) with a time step of 0.002 ps, was used to limit all of the bond lengths. In NVT and NPT ensembles, the medium’s complexes were able to equilibrate for 100 ps. MDS lasting 20 ns was run for each complex. For the analysis, all trajectories were saved every 2 ps.

### 2.5. Analysis of the MDS Results

The structural analysis of the mutant p53 and the selected phytocompound/s was performed by the different in-built functions of GROMACS that used the trajectory files as their inputs. The structural analysis comprised of the root mean square deviation (RMSD), root mean square fluctuation (RMSF), the radius of gyration (Rg), solvent accessible surface area (SASA), and the number of hydrogen bonds between the receptor and the ligands during the simulation, Graphical plots for the obtained trajectories were produced by the xmgrace program (Turner, 2005).

## 3. RESULTS

### 3.1. Comparative Modeling of the Mutant p53 and their Validation

The models of R175H, R248Q, and R273H mutant p53 were used for validation by the SAVES server. The programs of the SAVES server: PROCHECK, ERRAT, and VERIFY 3D resulted as follows. PROCHECK analysis provided a Ramachandran plot for the given protein structures (Modeller-generated P53WT, R175H, R248Q, and R273H models). The results of PROCHECK’s Ramachandran plot for all the models (wild-type (WT) and the mutant p53) are shown in Figure 1 and their statistics are enlisted in Table 1 and Supplementary Data S1. The Ramachandran plot shows that all the residues of P53WT, R175H, R248Q, and R273H lie in the various sub-regions of the allowed region (i.e. most favored region, additionally allowed region, and generously allowed region). Thus, all the residues fall within the allowed region.

**Table 1:** Residues of the generated mutant p53 models in the Ramachandran plot showing the quality of the modeled secondary structures.

| Regions of the Ramachandran Plot |  | P53WT | R175H | R248Q | R273H |
| --- | --- | --- | --- | --- | --- |
| Allowed region | Percentage of residues in the most favored region | 91.2 | 92.4 | 93 | 92.4 |
|  | Percentage of residues in additionally allowed region | 8.2 | 7.0 | 6.4 | 7 |
|  | Percentage of residues in generously allowed region | 0.6 | 0.6 | 0.6 | 0.6 |
| Disallowed region | Percentage of residues in disallowed region | 0 | 0 | 0 | 0 |
|  | Number of non-glycine and non-proline residues (%) | 100 | 100 | 100 | 100 |

**Figure 1.**
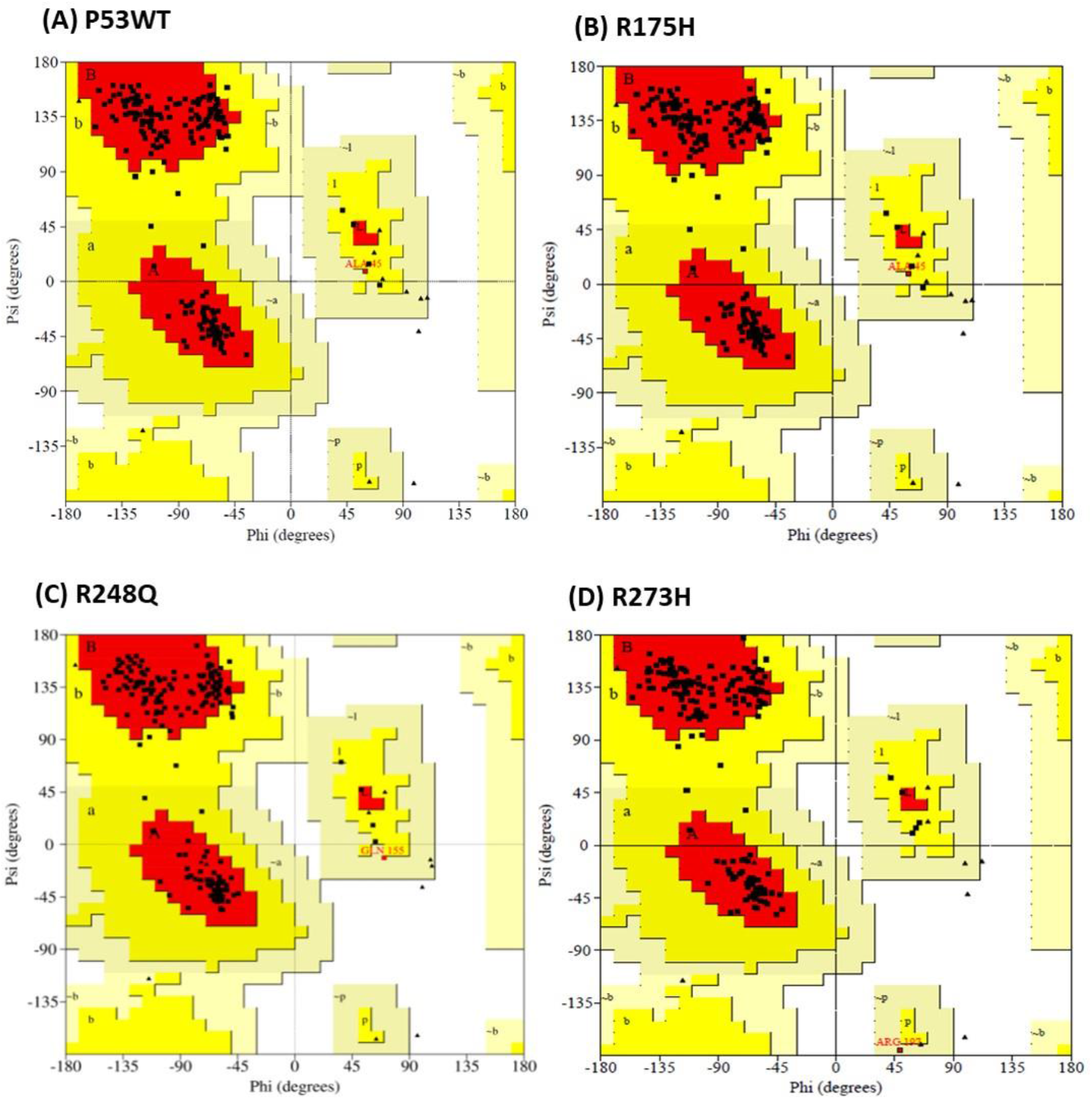
Ramachandran Plot obtained from the PROCHECK function analysis provided by SAVES online server

Table 2 and Supplementary Data S2 displays the validations performed by ERRAT and VERIFY 3D. The proportion of the protein for which the projected error is less than the 95% rejection level is the overall quality factor (OQF), which is provided by ERRAT. Furthermore, structures with an efficiently high resolution usually show an overall quality score of 95% or more. Low resolutions i.e. 2.5 to 3A usually produce an OQF of around 91%. The OQF projected by ERRAT for the P53WT, R175H, R248Q, and R273H models is 92.105%, 94.7917, 89.5833, and 92.1875, respectively. According to VERIFY3D results (Table 2), Modeller-generated P53WT, R175H, R248Q, and R273H models passed the structure validation with average scores of 93%, 90.5%, 99%, and 100% of their respective residues, which had average scorings of less than or equal to 0.2.

**Table 2:** Results of the structure validation by different functions of SAVES.

| Functions |  | P53WT | R175H | R248Q | R273H |
| --- | --- | --- | --- | --- | --- |
| ERRAT | Overall Quality Factor | 92.105 | 94.7917 | 91.1458 | 92.1875 |
| VERIFY3D | Status | Pass | Pass | Pass | Pass |
| | Residues having averaged 3D-1D score $\geq 0.2$ | 93% | 90.50% | 99% | 100% |
| | Amino acids having score $\geq 0.2$ in the 3D/1D profile | 80% | 80% | 80% | 80% |

The superimposition of generated P53WT, R175H, R248Q, and R273H models on the template further confirmed the deviation between the model and the template. As shown in Figure 2, the superimposition revealed an RMSD value of 0.212 Å, 0.240 Å, 0.231 Å, and 0.242 Å for P53WT, R175H, R248Q, and R273H, respectively.

**Figure 2.**
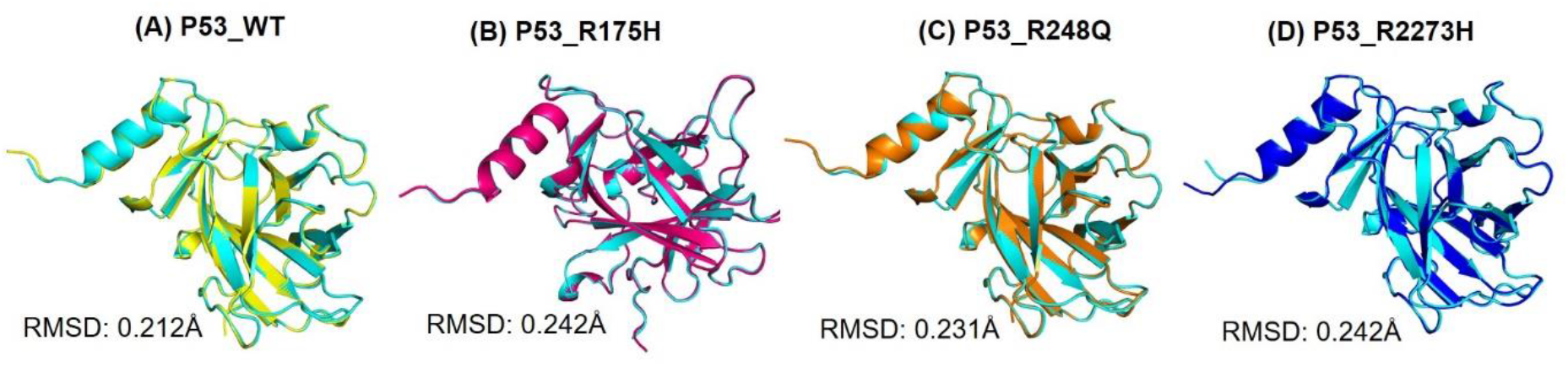
Superimposition of the Modeller generated P53WT, R175H, R248Q, and R273H models with the template (PDB ID:2AHI). The blue-colored structure indicates the template.

### 3.2. Molecular Docking with Autodock Vina

Validated models of R175H, R248Q, and R273H was considered as receptor for the molecular docking. Selected 60 compounds derived from AP and CA were subjected to docking at the defined drugable pocket of the R175H, R248Q, and R273H resulting in different confirmations of the ligand-receptor complex along with their respective binding energies. The Autodock Vina predicted binding energies of the first confirmations with zero RMSD values were selected. The binding energies of the docked results are enlisted in Supplementary Data S3, S4 and S5. We did not include any sets of controls in this investigation because, to our knowledge, no drugs were found to bind with the selected three mutants. Since there is no reference point to compare the binding energy of the docked phytocompounds, the top 5 compounds showing the minimum binding energies were considered for further study. However, for mutant R175H, a total of 6 phytocompounds were chosen because the binding energies of the 5^th^ and 6^th^ phytocompounds were identical. The binding energies of the docked mutants and the selected phytocompounds are shown in Figure 3. Repeats from the chosen phytocompounds were removed, leaving a total of 11 phytocompounds to be predicted to have drug-like properties.

**Figure 3.**
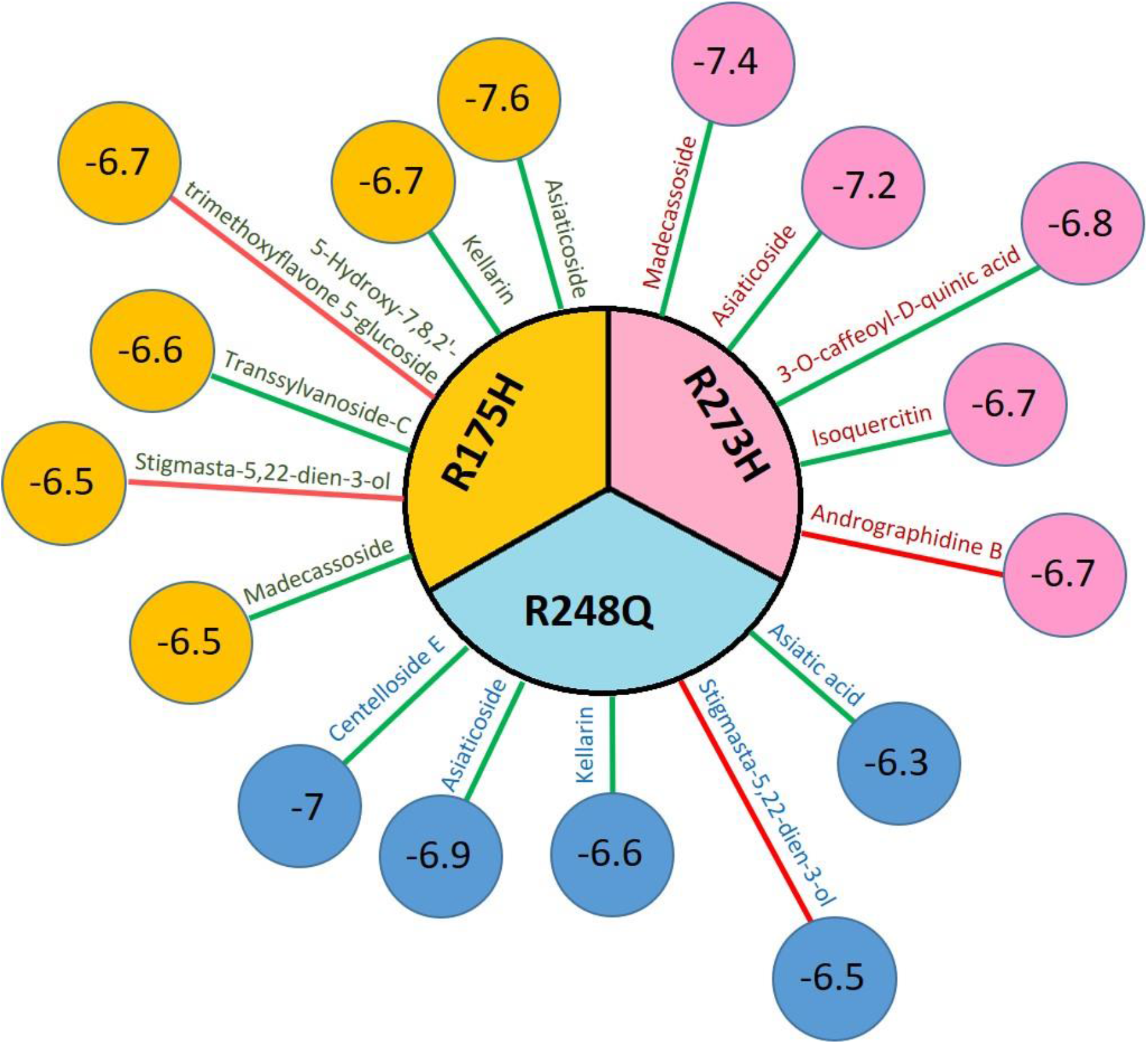
The phytocompounds selected based on their lower binding energies toward the receptors R175H, R248Q, and R273H. The edges represent the phytocompounds and the color of the edges represents the respective source plant for the phytocompounds (i.e. green: CA and red: AP. The small circles at the endpoint represent binding energies in Kcal/mol.

### 3.3. Prediction of Drug-like Properties and Bioactivity Scores

The selected 11 phytocompounds’ drug-likeness predictions offered information on their molecular features, including their MW and vol, miLogP, TPSA, natoms, nrotb, hydrogen acceptors, and donors. As indicated in Table 3, only one phytocompound, (i.e.asiatic acid) was able to pass the methodologies’ elimination criterion (mentioned in methods) for having drug-like characteristics. Figure 4 displays the bioactivity score of asiatic acid obtained from Molinspiration. Asiatic acid showed a 0.91 and 0.66 bioactivity score towards the nuclear receptor ligand and enzyme inhibitors respectively.

**Table 3.**
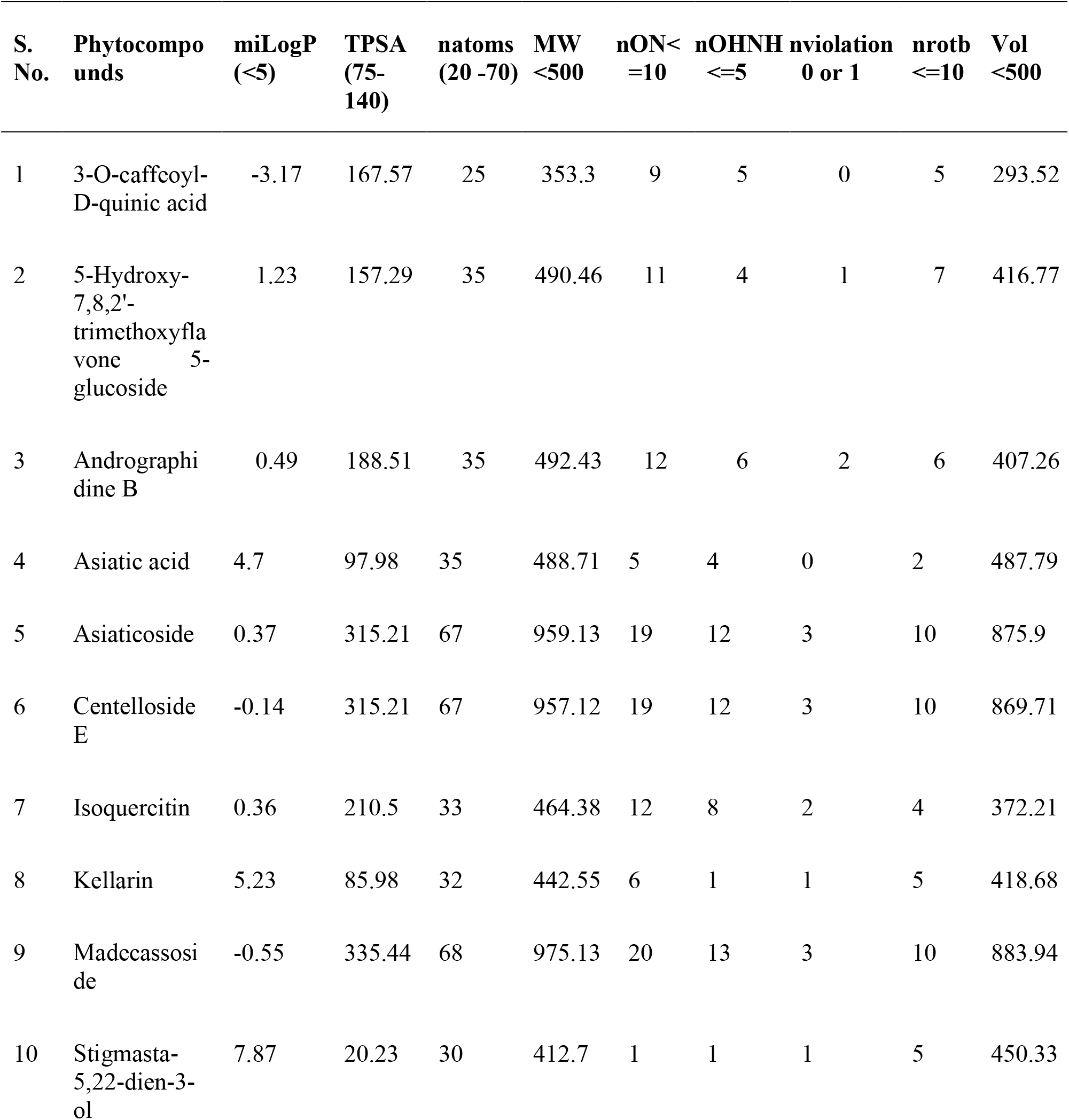
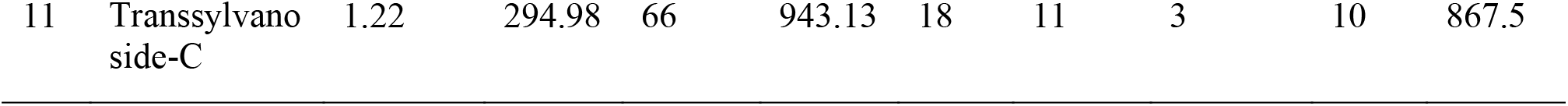
Molinspiration predicted drug likeness properties for the selected 11phytocompounds.

**Figure 4.**
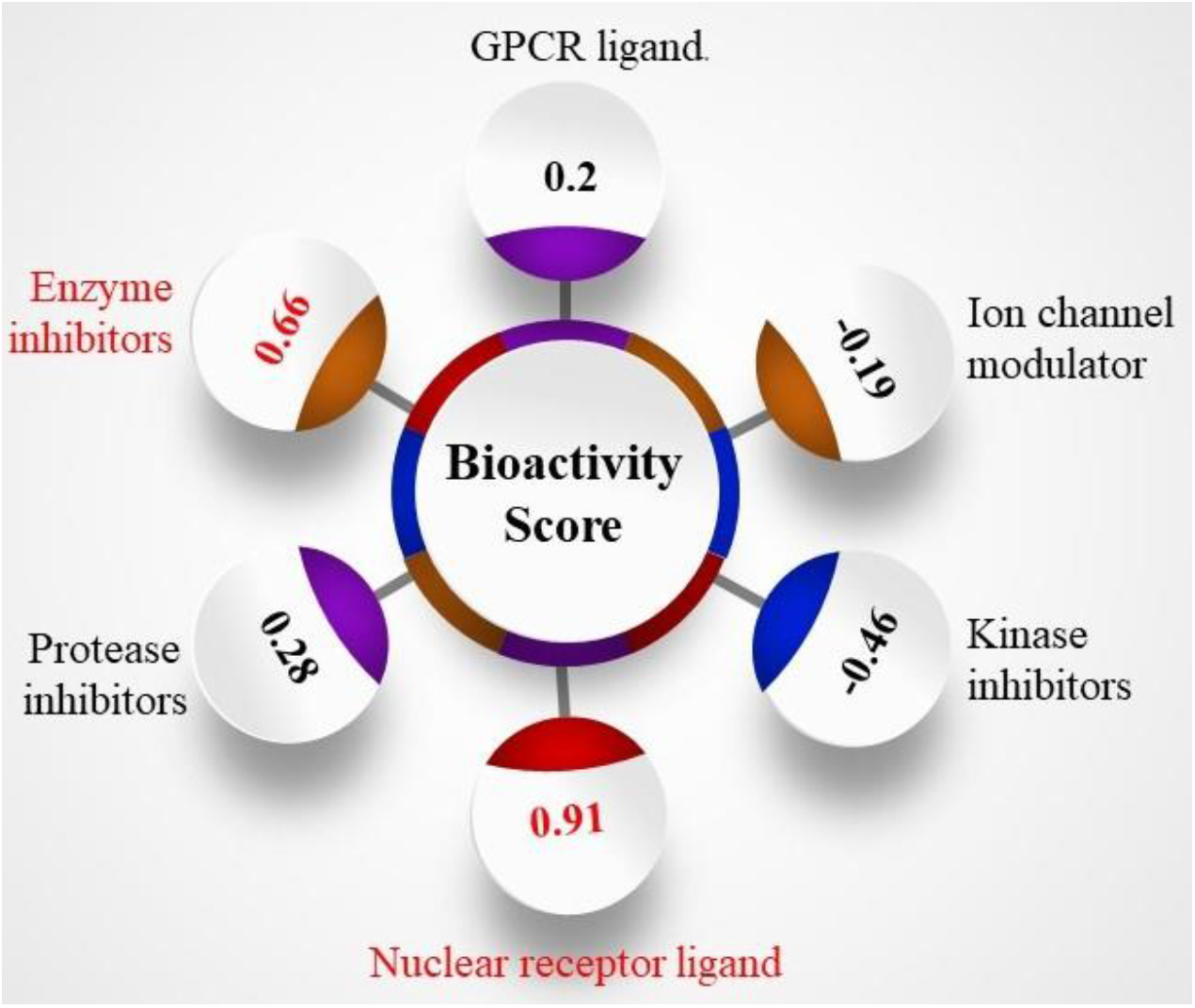
Bioactivity score of asiatic acid towards different targets, as calculated by Molinspiration. Asiatic acid showed the highest bioactivity score for the class of enzyme inhibitors and the nuclear receptor ligands.

### 3.4. Toxicity Prediction of Asiatic acid

Toxicity of asiatic acid predicted by ProTox-II server, an online tool that amalgamates 33 prediction models generated on the basis of *in vitro* as well as *in vivo* assay data. The SMILES notation of asiatic acid (CC1CCC2(CCC3(C(=CCC4C3(CCC5C4(CC(C(C5(C)CO)O)O)C)C)C2C1C)C)C(=O)O) obtained from the IMPPAT database was provided as input. ProTox-II offers information on acute toxicity (LD50, mg/kg), hepatotoxicity, and several toxicity endpoints such as carcinogenicity, mutagenicity, immunotoxicity and cytotoxicity to human cells. Figure 5 displays the predicted LD50 value, the class/category of toxicity, molecular similarity, and the prediction accurateness for asiatic acid. Taking into account the LD50 thresholds, the ProTox-II program creates six toxicity groups, ranging from the most hazardous to the least toxic. Here, the estimated median lethal dose (LD50) falls under the toxics of class 4 for acute oral toxicity (hazardous if swallowed), showing an LD50 value of 2000 mg/kg with an average molecular similarity of 89.49% and prediction accuracy of 70.97% (Figure 5). Asiatic acid was also identified as not being cytotoxic, mutagenic, carcinogenic, or poisonous to the liver. However, as can be seen in the radar chart (Figure 6), asiatic acid was also expected to have a small risk of immunotoxicity. Overall, as per the ProTox-II prediction, the asiatic acid displayed favourable toxicological profiles.

**Figure 5.**
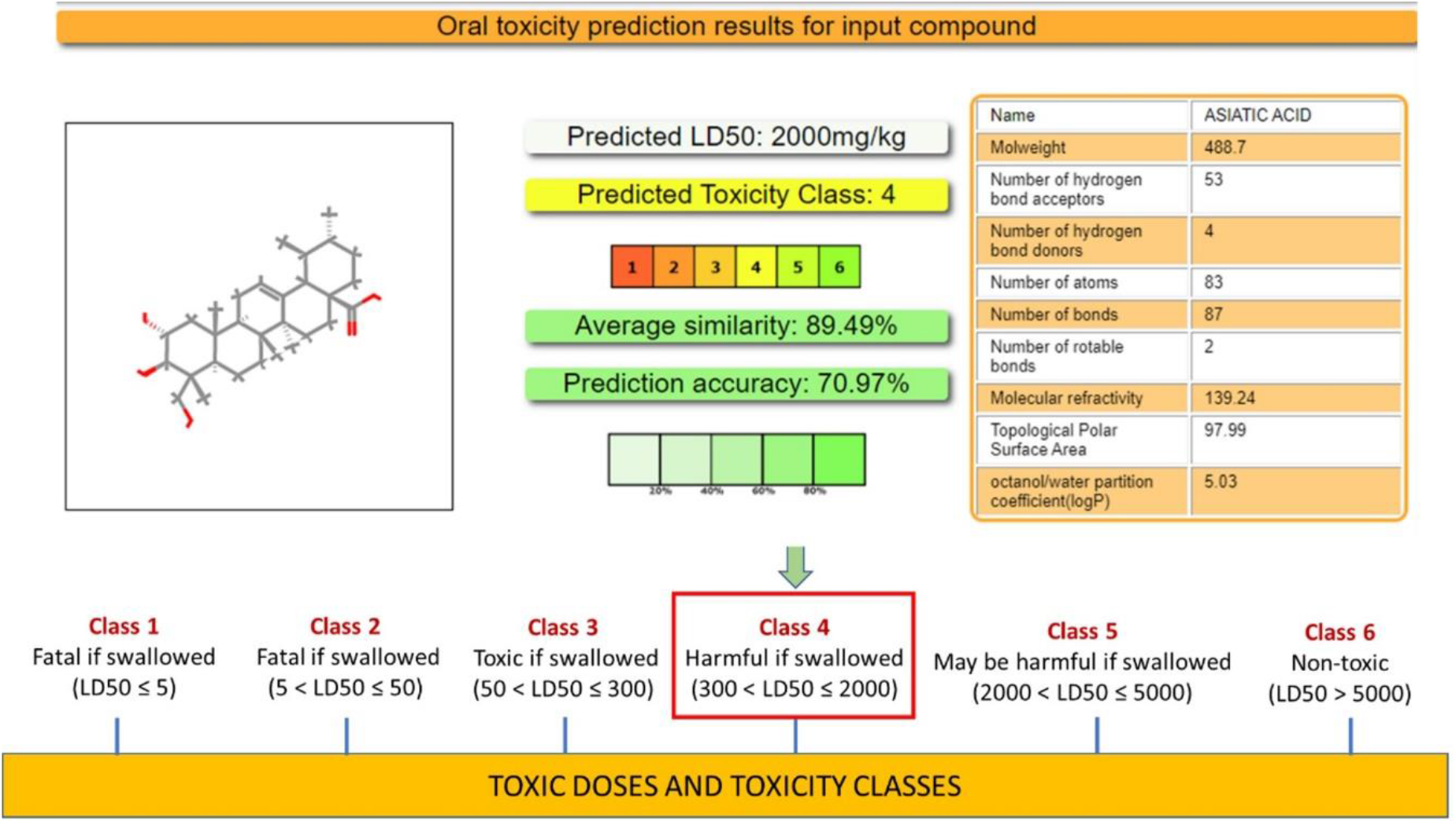
Oral toxicity prediction by ProTox-II. Asiatic acid falls under Class 4 toxicity.

**Figure 6.**
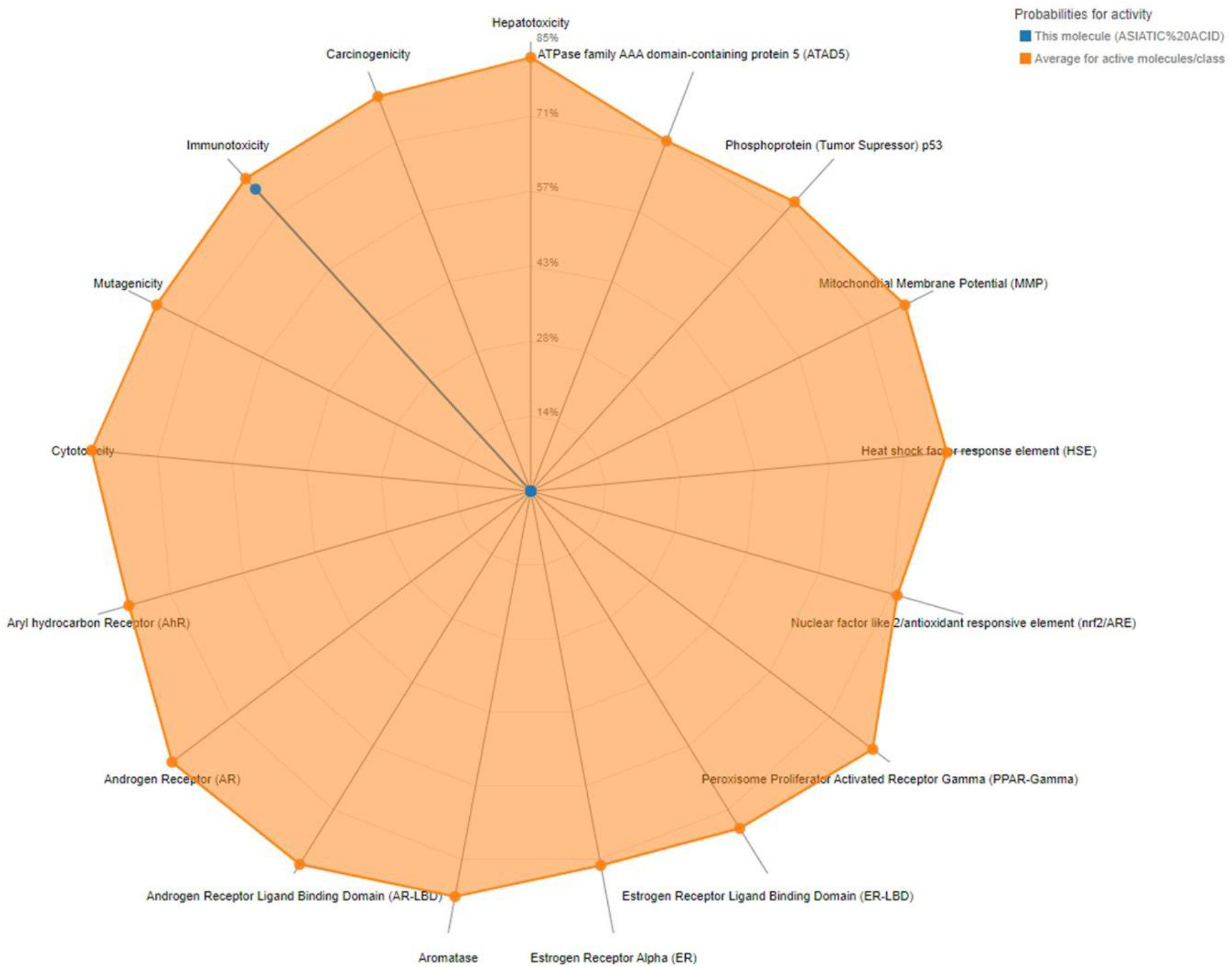
Toxicity radar charts for the predicted toxicity of asiatic acid. The chart indicates that asiatic acid is predicted to be active for one of the toxicity endpoints (i.e immunotoxicity).

### 3.5. Molecular Interactions of R248Q-Asiatic acid complex

The analysis of the interactions between R248Q and Asiatic acid after docking was visualized in Pymol and LigPlot+ shows that Asiatic acid may interact with R248Q via hydrogen bonds with the residue Gln-99 and Asp-114and hydrophobic bonds interactions with Pro-97, His-100, His-121, Arg-81 and Phe-119 as shown in Figure 7 (A) and (B).

**Figure 7.**
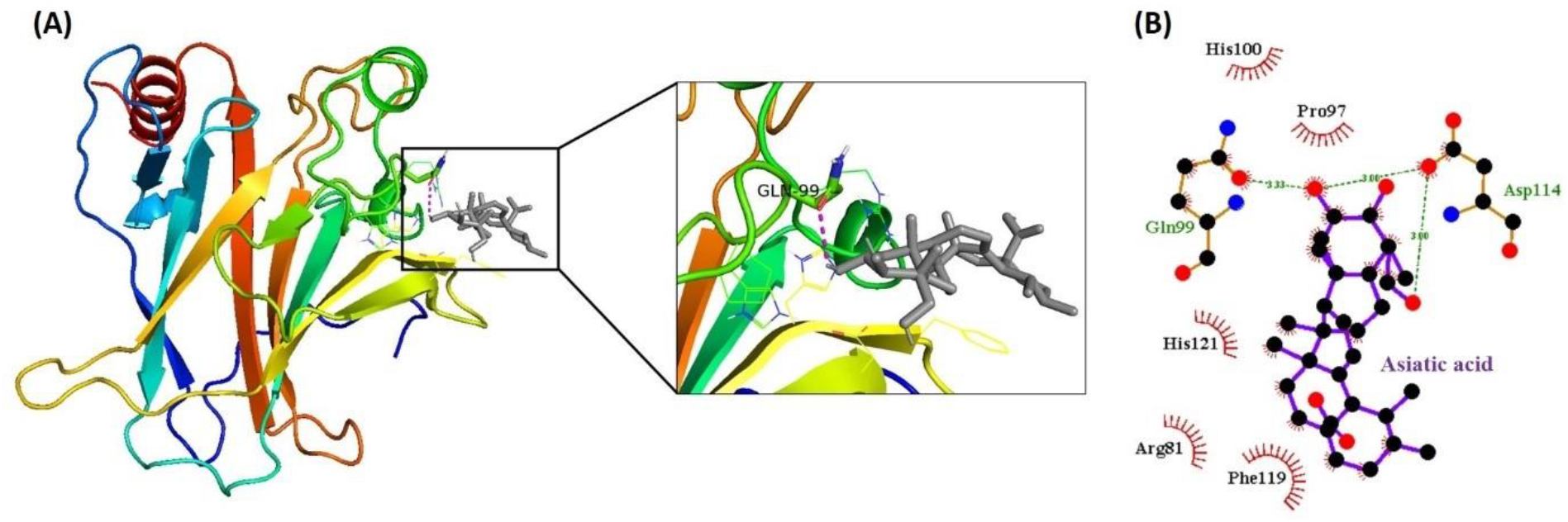
Molecular interactions of R248Q-Asiatic acid (A) hydrogen bonds visualized in PyMOL is denoted by the magenta dotted line and grey colored compound represents the asiatic acid. (B) hydrogen and hydrophobic bonds are visualized in LigPlot+, where the green dotted lines denote the hydrogen bonds and brown rays denote the hydrophobic interactions.

### 3.6. MDS of the R248Q-AA Docked Complex

MDS of 20 ns run resulted in trajectories for each system of wild-type p53 (P53WT), p53 mutant (R248Q), and R248Q-AA complex. The analysis of the RMSD, RMSF, Rg, SASA, as well as the hydrogen bond was done by generating their respective plots. The plots demonstrated a deviation, fluctuation, SASA, and stability of P53WT, R248Q, and R248Q-AA complexes throughout the MDS of 20 ns. Observed RMSD values showed that R248Q_receptor (green) has almost similar fluctuation to the P53WT (black) exhibiting an average RMSD of 0.2 nm. But after a 5ns simulation, the R248Q-AA complex (red) has higher fluctuation with the values ranging between 0.2-0.3 nm till the completion of MDS time period (i.e. 20 ns) as shown in Figure 8 (A).

**Figure 8.**
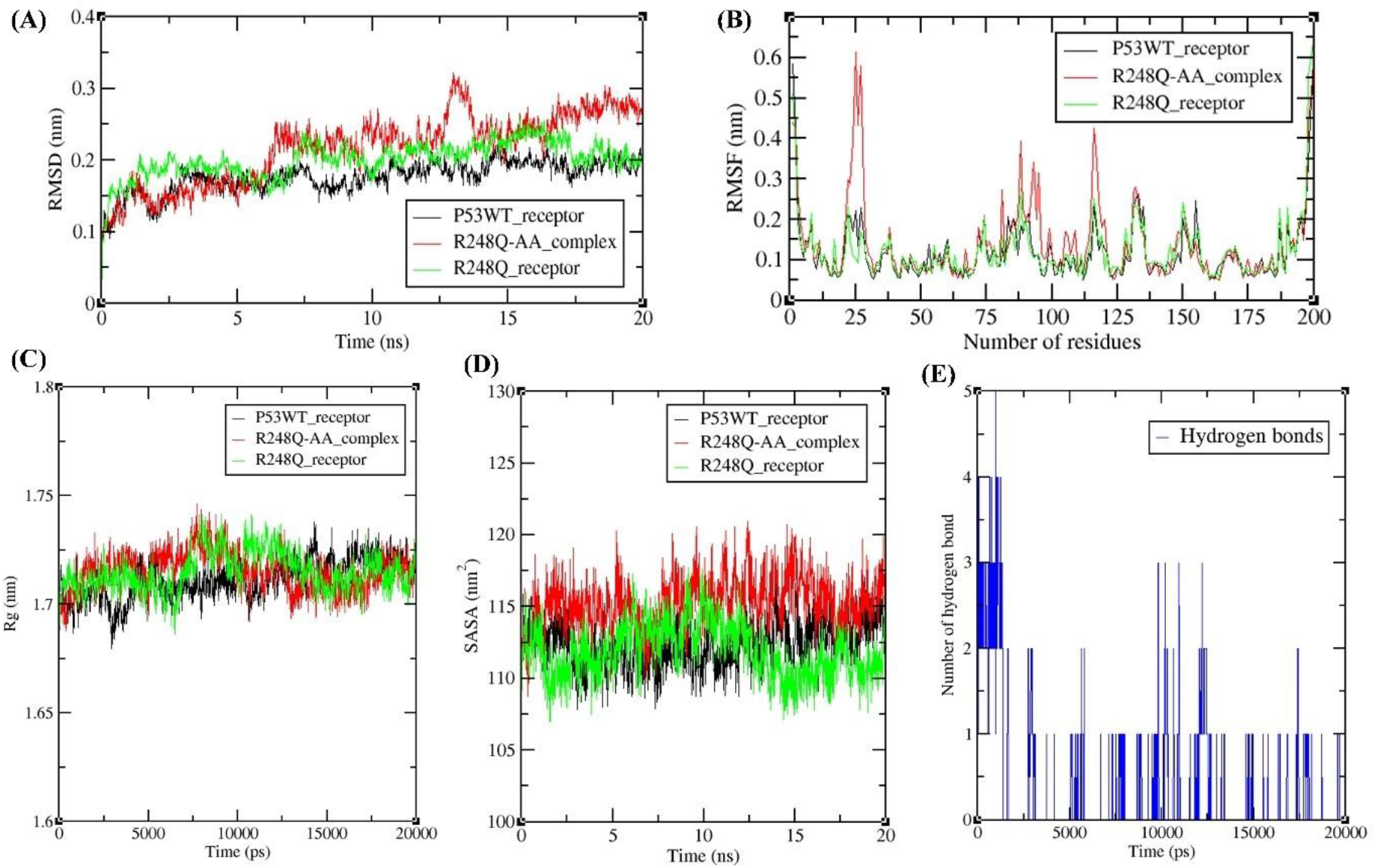
Graphical representation of the MDS analysis results (A) RMSD plot of P53WT receptor (Black color), R248Q-AA complex (red color), and R248Q receptor (green color) displaying the deviation and stability throughout the 20 ns MDS (B) The RMSF graph showing fluctuations per residue. (C) The Rg graph demonstrating the stability as well as the compactness of P53WT, R248Q, and R248Q-AA complex during 20,000 ps simulation (D) Plot showing the SASA of the P53WT, R248Q, and R248Q-AA complex during 20 ns simulation. (E) Hydrogen bond plot showing the hydrogen bonds formation between R248Q-AA throughout the 20 ns MDS.

The RMSF plot for all selected molecules showed an average fluctuation value of 0.1 nm as shown in Figure 8 (B). But some fluctuations as seen at 20-25, and 80-120 amino acid residue regions of R248Q-AA complex. It demonstrates that the residues at 20-25, and 80-120 regions are crucial for the interaction of R248Q-AA. The residues lying within these ranges are also observed in the hydrogen bond formations and hydrophobic interactions in the docking analysis as shown in Figure 6. Residues of the R248Q-AA complex have highly fluctuated in the AA binding site (i.e. Gln-99 and Asp-114).

Analysing the Rg is very significant for assessing the compactness as well as the stability of the protein structure throughout the MDS run time. Rg plot for all three molecules has shown Rg value ranged from 1.7-1.72 nm and an average of approximately 1.71 nm. Relatively invariant Rg values of P53WT in the plot show that the protein remains very stable, in its compact (folded) form throughout 20 ns. But the R248Q receptor and R248Q-AA complex showed a high Rg value at 8000 ps and later remains stable till the completion of 20000 ps. Overall, the Rg analysis suggested that P53WT, R248Q, and R248Q-AA maintained their stability as well as compactness for the 20ns MDS as shown in Figure 8(C).

SASA calculated for the resultant trajectories from MDS for every individual residue of P53WT, R248Q and R248Q-AA varied from 110-112 nm^2^, 108-112 nm^2^, and 112-118 nm^2^ respectively (Figure 8(D). Thus, the R248Q-AA has a relatively higher SASA as compared to the P53WT and R248Q receptors.

The hydrogen bond plot for R248Q-AA complex has shown the formation of 1-5 hydrogen bonds and AA interaction with R248Q formed an average of two hydrogen bonds during the 20 ns MDS as shown in Figure 8(E).

## 4. DISCUSSION

Warbug’s effect is a phenomenon, where the reliance of cancer cells on the energy shifts from OXPHOS to aerobic glycolysis. Thus, targeting the energy metabolism of cancer cells has attained importance as a crucial target in cancer therapeutics. Moreover, recent evidence has shown the association of mutated p53 with energy metabolism in cancer cells. More than 50 percent of cancers contain loss of function (LOF) mutations in *p53* (Ozaki and Nakagawara, 2011). Despite the existence of many mutations of p53, the hotspot mutations (R175H, R248Q, and R273H) are mainly reported for their contribution to the energy metabolism in cancer cells (Zhang et al., 2013; Chiang et al., 2021). Apparently, National Cancer Institute, GDC Data Portal reports the mutation R175H to be frequently occurring and exhibits their function via direct binding to promoters of genes and transcriptionally up- or down-regulating their expressions. For instance, R175H interacts with p63, p53, ETS2, and NF-Y to impede the target gene expressions. Other two mutants, R273H and R248Q were observed to perform similar functions as R175H. Under normal oxygen conditions, the p53 mutant R248Q favors aerobic glycolysis via upregulating expression of glycolytic enzymes but downregulating the proteins of OXPHOS. In a similar context, it is observed that the cancer cell carrying P53WT and OXPHOS-dependent ATP generation can be treated with mitochondria-targeting drugs. On the other hand, glycolytic inhibitors are suggested to be effective against R248Q-carrying as well as glycolysis-dependent cancer cells. Another study suggests that R175H and R273H mutations induce the citrate transport protein, important to interchange mitochondrial citrate and cytosolic malate for stimulating respiration as well as helping in maintaining the MIM integrity (Kolukula et al., 2014). In tumor cells, the mitochondrial citrate transporter gene preserves mitochondrial integrity, and bioenergetics from mitochondrial damage and depletion through autophagy, thus, encouraging proliferation (Catalina-Rodriguez, et al., 2012). Expression of the p53 mutants R175H, R248Q, and R273H in human lung carcinoma cells stimulates the Warburg effect, and thereby increases the level of glucose uptake, glycolytic rate as well as lactate production (Zhang et al., 2013). Under a condition of oxidative insult of cancer cells, R273H help in inhibiting the expression of detoxifying enzymes, such as heme oxygenase-1 (HO-1) as well as quinone oxidoreductase, thus facilitating the survival of these cells (Kalo et al., 2012). Thus, collectively, R175H, R248Q, and R273H seem to play an important role in improving the energy metabolism of cancer cells and thus may act as a significant target in cancer therapeutics.

Recent studies suggest the utilization of natural compounds to target different target of cancers due to their minimal side effects (Ali Abdalla et al., 2022; Lin et al., 2020; Hashem et al., 2022; Parate et al., 2021). In our recent research, the natural compounds from CA and AP were explored for their ability to target hexokinase 2 (overexpressed in the majority of the cancer cells and target for class I mitocans), Estrogen receptor, and Progesterone receptors (overexpressed in breast cancer cells) (Swargiary and Mani, 2021; Swargiary and Mani, 2022). By looking at the significance of the phytocompounds from both plants, as well as the published role of mutant p53 proteins in promoting the energy metabolism of cancer cells, it was further interesting to explore the ability of all these phytocompounds to target these p53 mutants. Initially, the molecular docking of 60 phytocompounds collectively from CA and AP was performed and was sorted based on their binding energies. As there were no control/s for comparing the binding energies, thus, the top 5 docked compounds were selected. However, in the case of the mutant R175H, 6 docked compounds were selected because the 5^th^ and 6^th^ compounds share the same binding energy. Thus, all the selected phytocompounds were segregated and only 11 phytocompounds remained after removing the repeats. For any compound to be called a drug should have drug-like properties, A traditional method to evaluate drug-likeness properties is to check whether the compound follows Lipinski’s Rule of Five that includes number of hydrophilic groups, MW, and hydrophobicity. However, in the current study, Lipinski’s rule, Veber’s rule, Ghosh’s rule, and Pfizer’s rule were combined for the selection of compounds, exhibiting drug-like properties. The parameters and their cutoffs were miLogP (<5), TPSA (75-140), natoms (20 -70), MW <500,nON<=10, nOHNH<=5, nviolation 0 or 1, nrotb<=10, Vol <500. Out of all the 11 compounds, only asiatic acid passed the rules of drug-likeness prediction. The bioactivity score of asiatic acid indicated it to behave as a good ligand against nuclear receptor (transcription factors) as well as enzyme inhibitor too. Interestingly the selected mutant p53 proteins still harbour the function of transcription factor (GOF), further supporting these mutants to behave as good target for asiatic acid. The toxicity analysis results suggest that asiatic acid showed no sign of being cytotoxic, mutagenic, carcinogenic, or poisonous to the liver. Although a small risk of immunotoxicity is excepted, but the overall observations demonstrate a favorable toxicological profile of asiatic acid in accordance with the ProTox-II server. Thus, the overall analysis from the molecular docking, drug-likeness properties, bioactivity score, and toxicity prediction shows that asiatic acid is the most important phytocompounds from all our selected 60 phytocompounds. From the docking results, the asiatic acid has revealed the greatest binding energy (−6.3 Kcal/mol) towards the mutant R248Q. Therefore, the R248Q-asiatic acid complex was subjected to MDS and to understand the changes upon exposure to the ligand, the results were compared with the MDS for wild-type p53, and the mutant R248Q. It was observed that the asiatic acid significantly interacts with the mutant R248Q via hydrogen bonds indicating that asiatic acid could be a prospective lead compound against the mutant R248Q.

As per the literature, any compounds that targeting the cancer cell’s mitochondria are called mitocans. Based on their targets/ modes of action they have been classified into different classes such as 1. Hexokinase inhibitors, 2. BH3 mimetics, and associated agents (those damaging the role of the anti-apoptotic Bcl-2 family proteins), 3. inhibitors of Thiol redox, 4. voltage-dependent anion channel (VDAC) and adenine nucleotide translocase (ANT) targeting agents, 5. respiratory chain complex (es) targeting compounds, 6. MIM targeting hydrophobic cationic compounds, 7. other compounds affecting the Kreb’s cycle, and 8. agents hampering the mtDNA. So, based on the definition of mitocans, we can propose that asiatic acid may behave as a potential mitocan by indirectly targeting the mitochondria of the cancer cell. Interestingly, the results of our study may also help in proposing a new target for mitocan i.e. mutant p53 targeting mitocans.

In support of our findings, the anticancer potential of asiatic acid on different cancer models have been reported (Chen et al., 2023; Hao et al., 2018; Li et al., 2021; Ren et al., 2016). Asiatic acid have also shown indirect inhibition of the P53WT expression.

Asiatic acid has been reported for induction of apoptosis via release of intracellular Ca2+ and enhanced p53 expression in human hepatoma cells (Lee et al., 2002) and inhibit the expression of p53 in human breast cancer cells (Gou et al., 2020). However, to our knowledge, the effect of asiatic acid on mutant p53 is not yet explored. As a result of our initial research, we are able to report for the first time that mutant p53 (particularly, R248Q) may be targeted by asiatic acid and that this possibility may be investigated in R248Q mutant models of the cancerous system in the future. Moreover, on the basis of different *in-silico* analyses, our previous study has suggested asiatic acid as a mitocan that can target hexokinase 2, which catalyzes the very first step in glucose metabolism (Swargiary and Mani, 2021). The study proposed that the inhibition of hexokinase 2 by asiatic acid may dissociate hexokinase 2 from VDAC leading to suppression of anti-apoptotic effect of the HK2-VDAC complex. Thus, the dissociation may cause the opening of VDAC, which results in cytochrome c release, thereby causing apoptosis of the cancer cells (Swargiary and Mani 2021). On the other hand, the current study suggests that asiatic acid may target the mutant p53 (R248Q) and due to this, the R248Q mutant may not exhibit its beneficial effects to the cancer cells, carrying this mutation. As asiatic acid is predicted here to lower the pro-tumor effects of mutant p53, hence it may be suggested to be used in combination with standard therapy/drug. Combining our previous and current study, we may propose a holistic effect of asiatic acid by targeting hexokinase 2 and mutant p53 (R248Q) that may, directly and indirectly, enable the targeting of energy metabolism in the cancer cells.

## 5. CONCLUSION

From this study, we conclude that the mutant p53 (R175H, R248Q, and R273H) contributing to the metabolic reprogramming of the cancer cells, can serve as a potential target for mitocans (Figure 9). Although there are different targets for mitocans, our study for the first time suggests mutant p53 as the potential target for the mitocans. From this *in-silico* study, we suggest that asiatic acid, the phytocompound of CA, has drug-like properties and may be a potential lead compound for targeting the p53 mutant R248Q. Further experimental validation is proposed to strengthen the findings of our study. In the future, efficient delivery of asiatic acid to the site of action may be explored. Moreover, the immunotoxicity of asiatic acid predicted by Protox-II suggests exploring ways to reduce the immunotoxic effect of asiatic acid.

**Figure 9.**
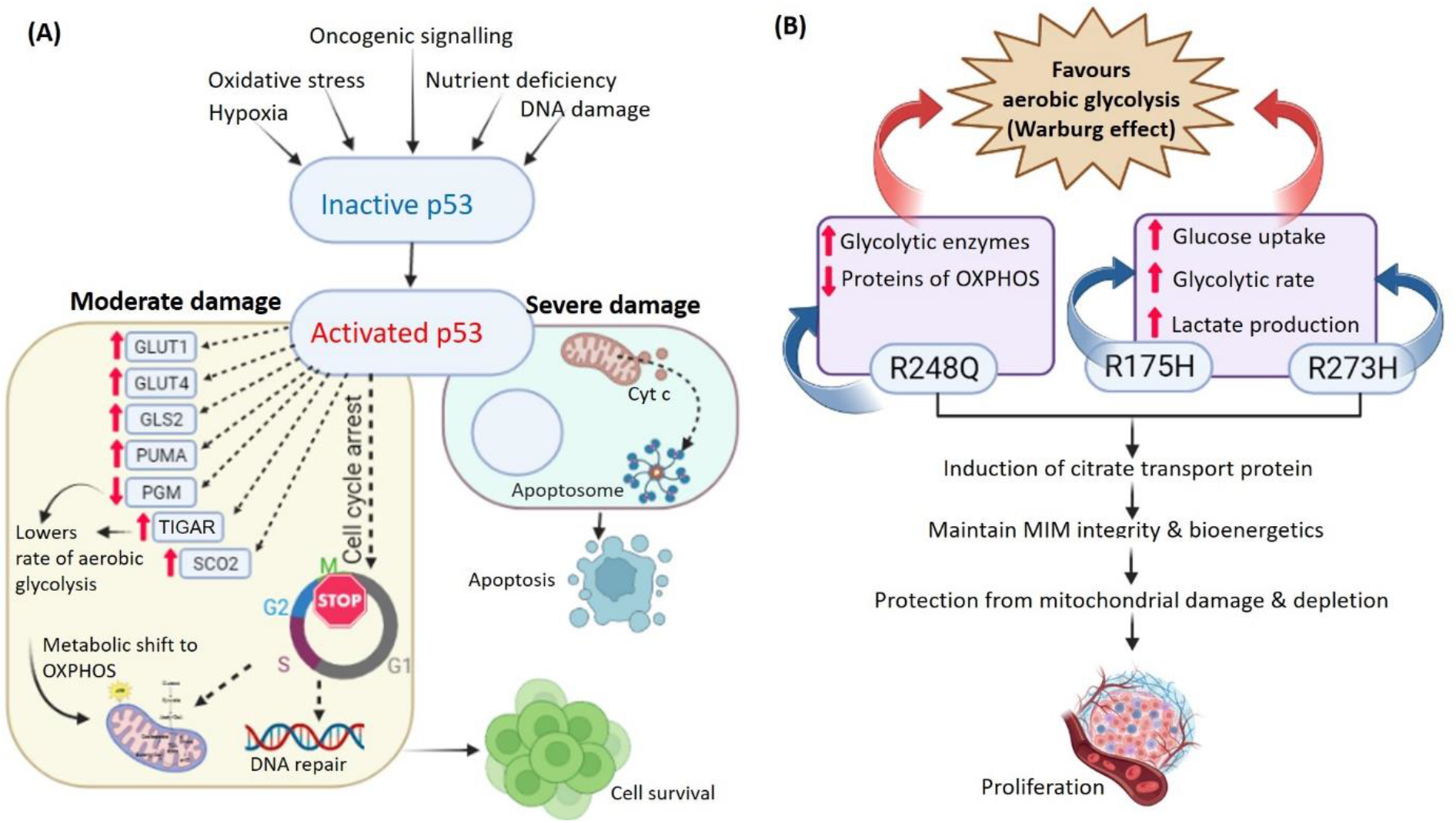
Showing the role of WT and mutant p53 in normal and cancer cells respectively. (A) In normal cell, the activated p53 under moderate stress conditions, help in DNA repairing and metabolic shifting of the cell, leading to cell survival. But in the severely damaged condition, the activated p53 triggers the cytochrome c release from mitochondria that consequently lead to apoptosis of the cell. (B) In cancer cell, the mutant p53 gets reactivated and triggers the expression of glucose transportes, OXPHOS proteins and citrate transport proteins, leading to increase in aerobic glycolysis and thus Warburg effects in cancer cells. Asiatic acid is proposed to target the R248Q, hence may affect the level of glycolytic enzymes and OXPHOS proteins in these cells.

## Supporting information

Supplementary Data S1. Ramachandran plot for the models generated by Modeller

Supplementary Data S2. Results from ERRAT and VERIFY 3D validation

Supplementary Data S3: Binding energies for docking of p53 mutation R175H

Supplementary Table S4: Binding energies for docking of p53 mutation R248Q

Supplementary Table S5: Binding energies for docking of p53 mutation R273H

## ACKNOWLEDGMENTS

The authors acknowledge Jaypee Institute of Information Technology, Noida, for providing the suitable infrastructure to complete this project. The authors are also grateful to the Ministry of Tribal Affairs, India.

