## Supplementary Data S1. Ramachandran plot for the models generated by Modeller for "Natural Product Screen Identifies Asiatic Acid Targeting Mutant p53"

**(A) P53WT model**

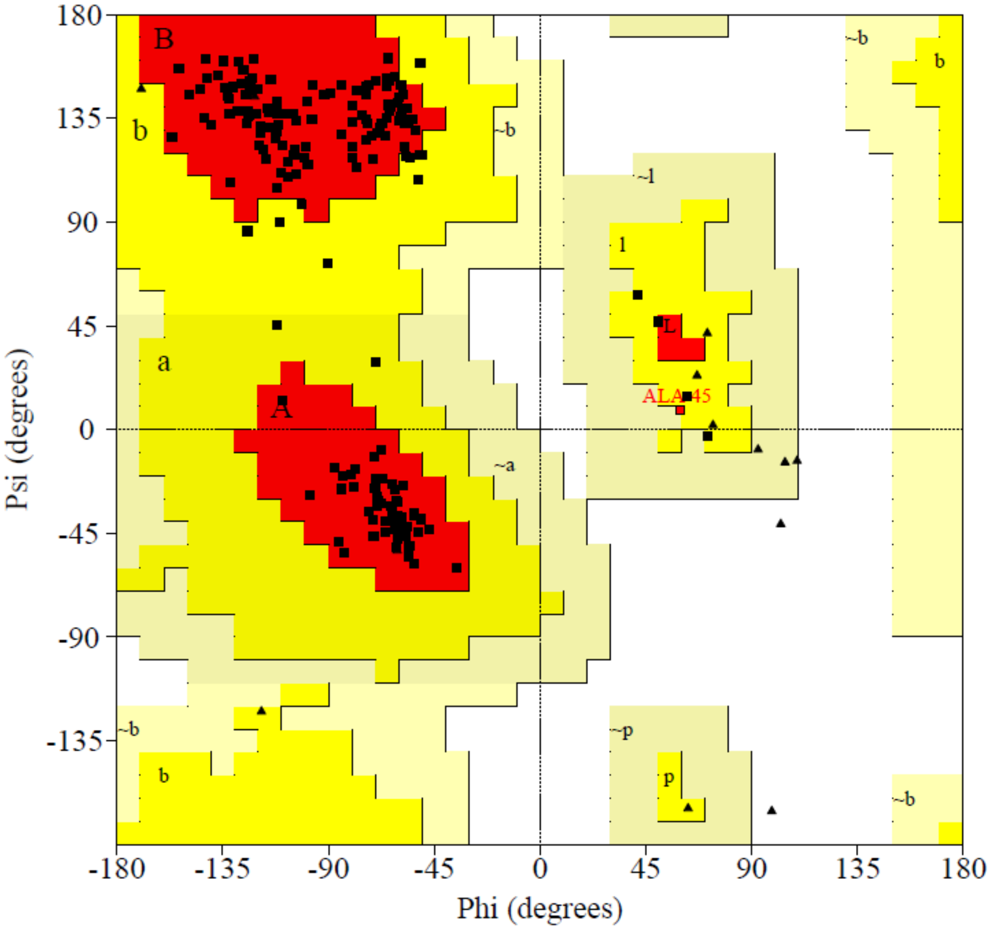


### Plot statistics

| | | |
|---|---|---|
| Residues in most favoured regions [A,B,L] | 156 | 91.2% |
| Residues in additional allowed regions [a,b,l,p] | 14 | 8.2% |
| Residues in generously allowed regions [~a,~b,~l,~p] | 1 | 0.6% |
| Residues in disallowed regions | 0 | 0.0% |
|  | ---- | ------ |
| Number of non-glycine and non-proline residues | 171 | 100.0% |
| Number of end-residues (excl. Gly and Pro) | 1 | |
| Number of glycine residues (shown as triangles) | 14 | |
| Number of proline residues | 14 | |
|  | ---- |  |
| Total number of residues | 200 | |

Based on an analysis of 118 structures of resolution of at least 2.0 Angstroms and R-factor no greater than 20%, a good quality model would be expected to have over 90% in the most favoured regions.

**(B) R175H model**

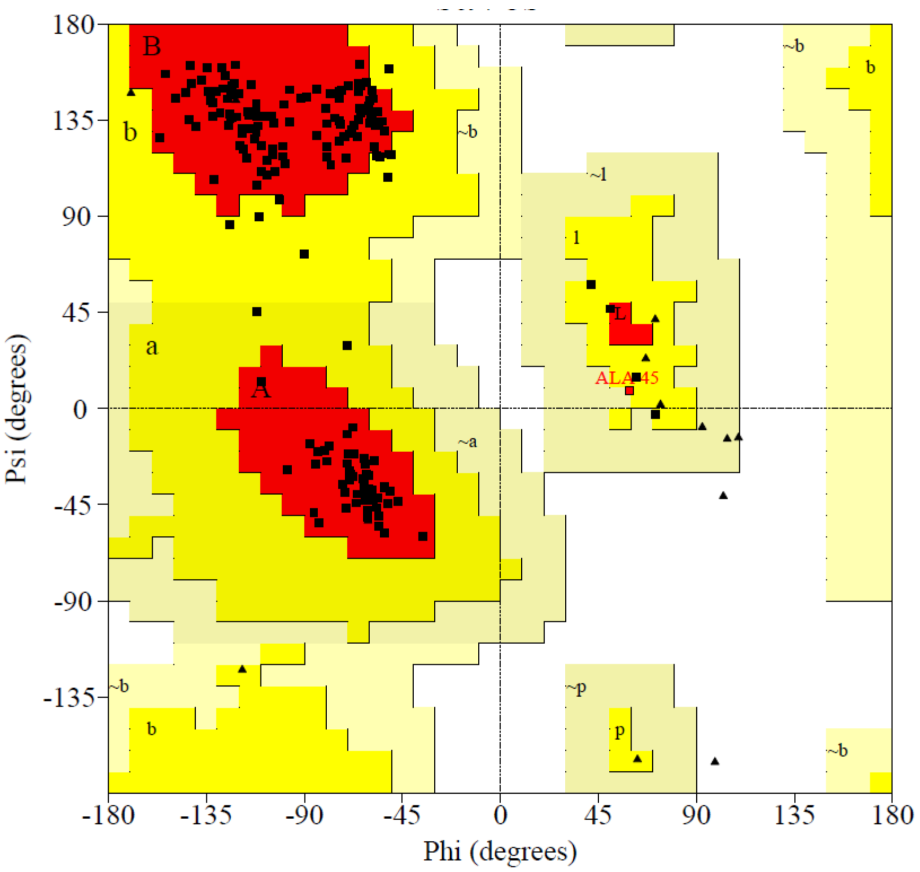


### Plot statistics

| | | |
|---|---|---|
| Residues in most favoured regions [A,B,L] | 156 | 91.2% |
| Residues in additional allowed regions [a,b,l,p] | 14 | 8.2% |
| Residues in generously allowed regions [~a,~b,~l,~p] | 1 | 0.6% |
| Residues in disallowed regions | 0 | 0.0% |
|  | ---- | ------ |
| Number of non-glycine and non-proline residues | 171 | 100.0% |
| Number of end-residues (excl. Gly and Pro) | 1 | |
| Number of glycine residues (shown as triangles) | 14 | |
| Number of proline residues | 14 | |
|  | ---- |  |
| Total number of residues | 200 | |

Based on an analysis of 118 structures of resolution of at least 2.0 Angstroms and R-factor no greater than 20%, a good quality model would be expected to have over 90% in the most favoured regions.

**(C) R248Q model**

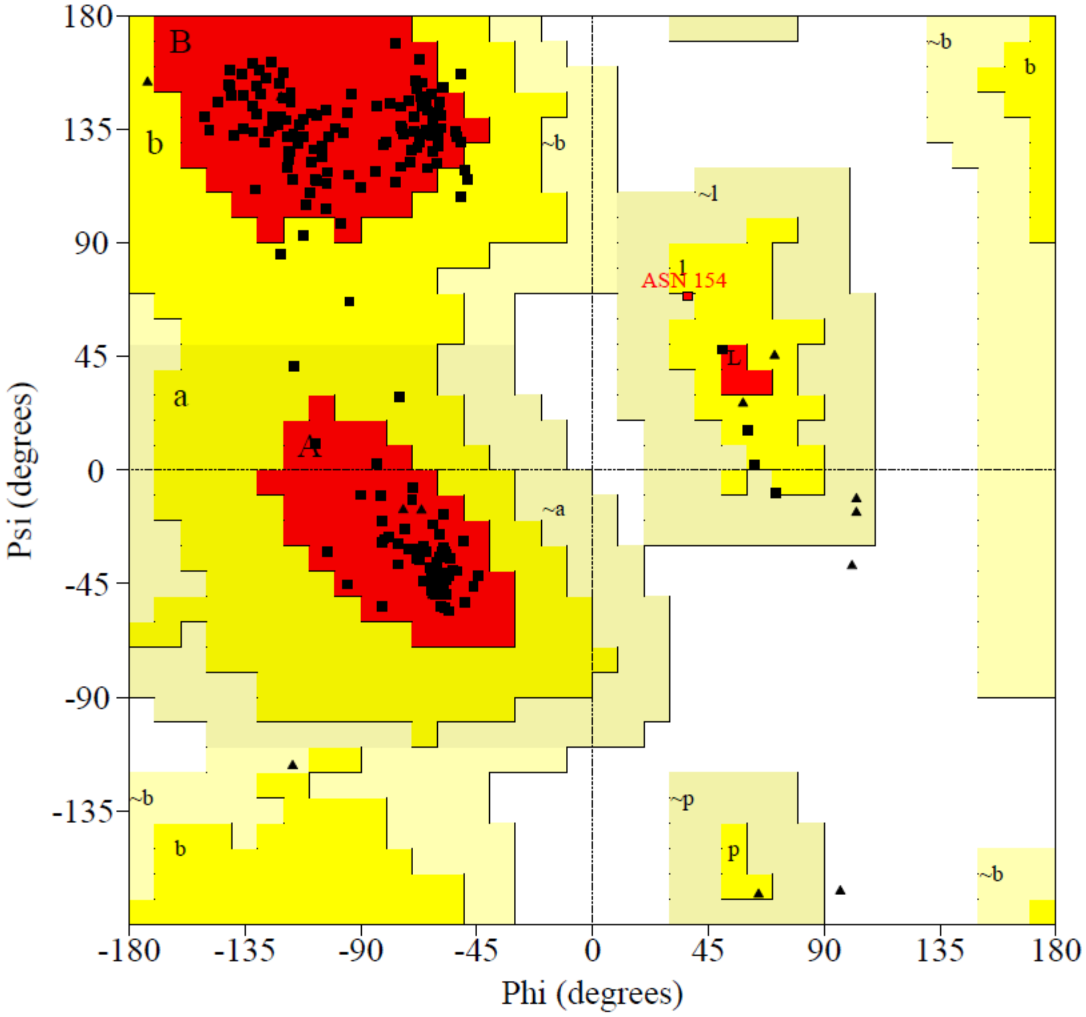


### Plot statistics

| | | |
|---|---|---|
| Residues in most favoured regions [A,B,L] | 159 | 93.0% |
| Residues in additional allowed regions [a,b,l,p] | 11 | 6.4% |
| Residues in generously allowed regions [~a,~b,~l,~p] | 1 | 0.6% |
| Residues in disallowed regions | 0 | 0.0% |
|  | ---- | ------ |
| Number of non-glycine and non-proline residues | 171 | 100.0% |
| Number of end-residues (excl. Gly and Pro) | 1 | |
| Number of glycine residues (shown as triangles) | 14 | |
| Number of proline residues | 14 | |
|  | ---- |  |
| Total number of residues | 200 | |

Based on an analysis of 118 structures of resolution of at least 2.0 Angstroms and R-factor no greater than 20%, a good quality model would be expected to have over 90% in the most favoured regions.

**(D) R273H model**

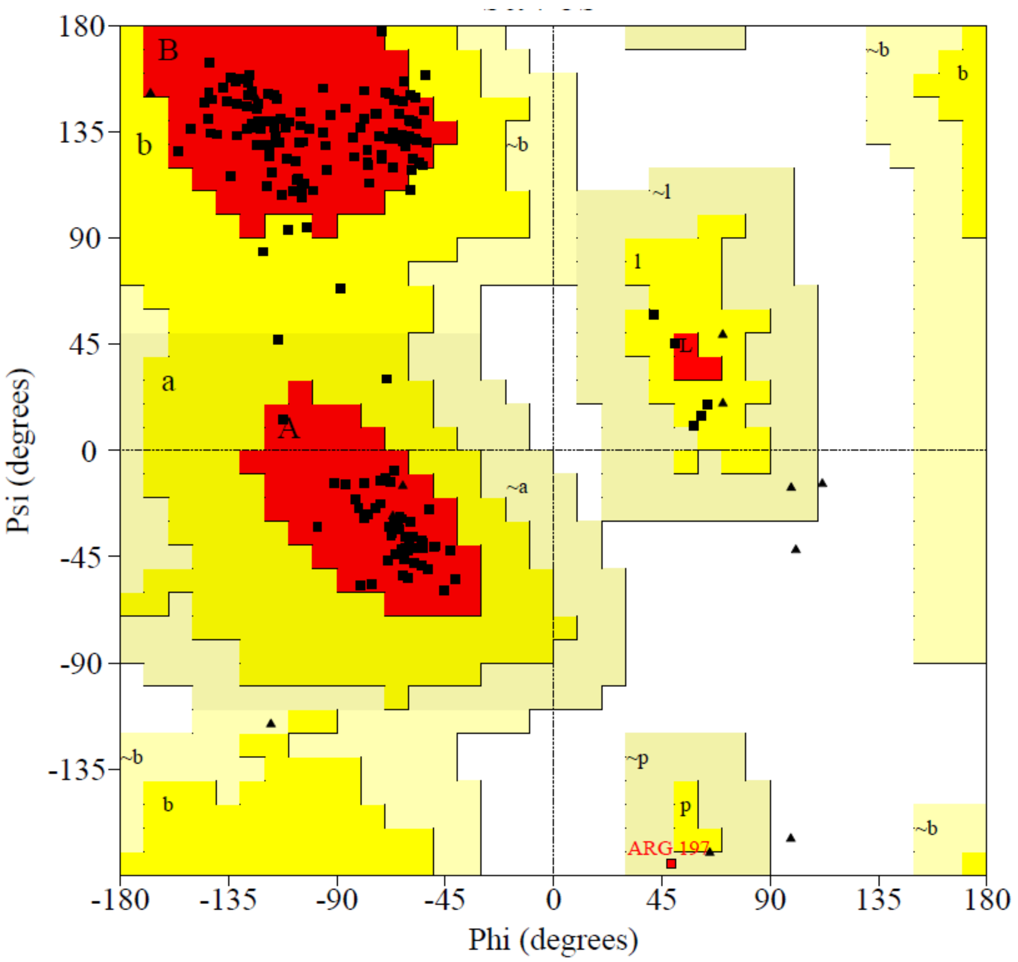


### Plot statistics

| | | |
|---|---|---|
| Residues in most favoured regions [A,B,L] | 158 | 92.4% |
| Residues in additional allowed regions [a,b,l,p] | 12 | 7.0% |
| Residues in generously allowed regions [~a,~b,~l,~p] | 1 | 0.6% |
| Residues in disallowed regions | 0 | 0.0% |
|  | ---- | ------ |
| Number of non-glycine and non-proline residues | 171 | 100.0% |
| Number of end-residues (excl. Gly and Pro) | 1 | |
| Number of glycine residues (shown as triangles) | 14 | |
| Number of proline residues | 14 | |
|  | ---- |  |
| Total number of residues | 200 | |

Based on an analysis of 118 structures of resolution of at least 2.0 Angstroms and R-factor no greater than 20%, a good quality model would be expected to have over 90% in the most favoured regions.
