## Supplementary figures and images for "Natural Product Screen Identifies Asiatic Acid Targeting Mutant p53"

### Supplementary Data S2. Results from ERRAT and VERIFY 3D validation

**Supplementary Data S2. Results from ERRAT and VERIFY 3D validation**

**(A) Wild type P53**

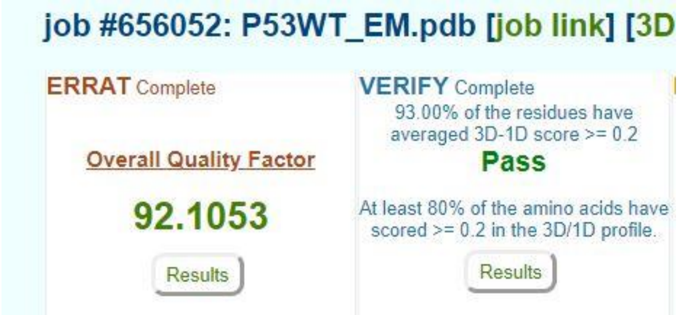


**(B) R175H**

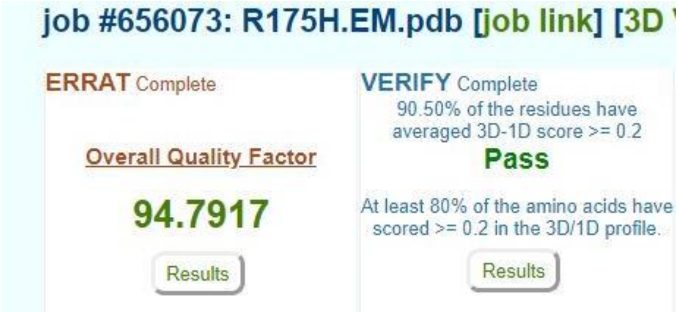


**(C) R248Q**

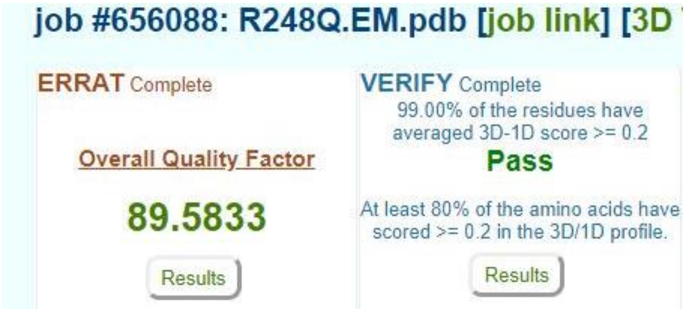


**(D) R273H**

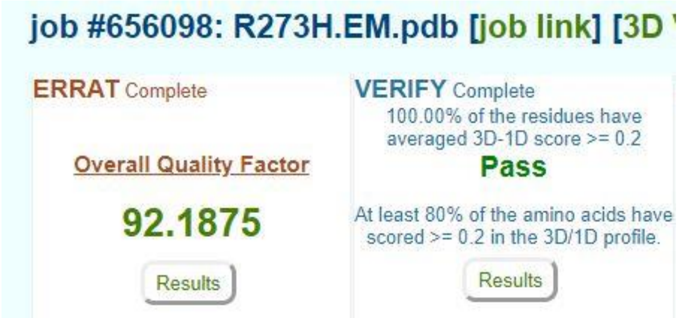
