## Supplementary Data S3: Binding energies for docking of p53 mutation R175H for "Natural Product Screen Identifies Asiatic Acid Targeting Mutant p53"

| S. No. | Plant | Compounds | Binding affinity (Kcal/mol) |
|---|---|---|---|
| 1 | CA | Asiaticoside | -7.6 |
| 2 | CA | Kellarin | -6.7 |
| 3 | AP | 5-Hydroxy-7,8,2'-trimethoxyflavone 5-glucoside | -6.7 |
| 4 | CA | Transsylvanoside-C | -6.6 |
| 5 | AP | Stigmasta-5,22-dien-3-ol | -6.5 |
| 6 | CA | Madecassoside | -6.5 |
| 7 | CA | isoquercitin | -6.4 |
| 8 | CA | Centelloside E | -6.4 |
| 9 | AP | Andrographidine B | -6.4 |
| 10 | AP | 14-Deoxyandrographoside | -6.3 |
| 11 | AP | Andrographiside | -6.3 |
| 12 | AP | Citrostadienol | -6.2 |
| 13 | CA | bayogenin | -6.2 |
| 14 | AP | Andrographidine E | -6.1 |
| 15 | CA | Hyaluronic acid | -6.1 |
| 16 | CA | Phytosterols | -6 |
| 17 | AP | Andrographidine D | -6 |
| 18 | CA | Asiatic acid | -5.9 |
| 19 | CA | AC1L4F91 | -5.9 |
| 20 | AP | Andrographidine C | -5.9 |
| 21 | AP | Andrographidine A | -5.9 |
| 22 | AP | Andropanoside | -5.9 |
| 23 | AP | Diterpene II (Lactone) | -5.8 |
| 24 | CA + AP | 3-O-caffeoyl-D-quinic acid | -5.8 |
| 25 | CA | Ginsenosides | -5.8 |
| 26 | CA + AP | Apigenin | -5.8 |
| 27 | CA | Dammarane | -5.8 |
| 28 | CA | Isochlorogenic acid | -5.8 |
| 29 | CA | farnesyl diphosphate | -5.7 |
| 30 | CA | Astragalin | -5.7 |
| 31 | AP | 14-Deoxy-11,12-didehydroandrographolide | -5.7 |
| 32 | CA | kaempferol | -5.6 |
| 33 | AP | Deoxyandrographolide | -5.6 |
| 34 | CA | Anthrone | -5.5 |
| 35 | AP | Andrographidine F | -5.5 |
| 36 | AP | Andrographin | -5.4 |
| 37 | AP | 14-deoxyandrographolide | -5.4 |
| 38 | AP | 5-Hydroxy-3,7,8-trimethoxy-2-(2-methoxyphenyl)-4H-chromen-4-one | -5.4 |
| 39 | AP | 5-hydroxy-7,8,2',3'-tetramethoxyflavone | -5.4 |
| 40 | AP | Dehydroandrographoline | -5.3 |
| 41 | AP | Andrographolide | -5.3 |
| 42 | AP | Andrograpanin | -5.2 |
| 43 | AP | 14-Deoxy-11-oxoandrographolide | -5.1 |
| 44 | CA | alpha-Chamigrene | -5 |

| | | | |
|---|---|---|---|
| 45 | AP | MLS001143515 | -5 |
| 46 | CA | Hydrocotoin | -4.9 |
| 47 | CA | 4-aminobutyric acid | -4.8 |
| 48 | CA | UNII-0V56HXQ8N5 | -4.7 |
| 49 | CA | l-ascorbic acid | -4.6 |
| 50 | CA | ferulic acid | -4.5 |
| 51 | AP | 2,4-Dihydroxycinnamic acid | -4.5 |
| 52 | AP | CARVACROL | -4.4 |
| 53 | CA | alpha-Terpineol | -4.4 |
| 54 | CA | Gulonic acid | -4.4 |
| 55 | CA | DL-Alanine-15N | -4.3 |
| 56 | CA | alpha-Humulene | -4.3 |
| 57 | CA | 4-Methoxybenzaldehyde | -3.9 |
| 58 | CA | 2,3-Dihydrobenzofuran | -3.8 |
| 59 | CA | 3-Carene | -3.7 |
| 60 | CA | 2-Methyl-2-Butanol | -3.1 |
